# Functional connectivity reveals dissociable ventrolateral prefrontal mechanisms for the control of multilingual word retrieval

**DOI:** 10.1101/354829

**Authors:** Francesca M. Branzi, Clara D. Martin, Manuel Carreiras, Pedro M. Paz-alonso

## Abstract

A fundamental cognitive operation involved in speech production is word retrieval from the mental lexicon, which in monolinguals is supported by dissociable ventro-lateral prefrontal cortex (vlPFC) mechanisms associated with proactive and reactive control. This functional magnetic resonance imaging (fMRI) study established whether in multilinguals word retrieval is supported by the same prefrontal mechanisms, and whether proactive modulation consists in suppression of the non-target lexicon. Healthy multilingual volunteers participated in a task that required them to name pictures alternatively in their dominant and less-dominant language. Two crucial variables were manipulated: the *cue-target interval* (CTI) to either engage (long CTI) or prevent proactive control processes (short CTI), and the *cognate status* of the pictures to-be-named (non-cognates *versus* cognates) to capture the presence of selective pre-activation of the target language. Results support the two-process account of vlPFC and indicate that multilinguals engage in proactive control to prepare the target language. This proactive modulation, enacted by anterior vlPFC, is achieved by boosting the activation of lexical representations of the target language. Control processes supported by mid-vlPFC and left inferior parietal lobe together, are similarly engaged in pre-and post-word retrieval, possibly exerted on phonological representations to reduce cross-language interference.

Significance Statement

Word retrieval in monolingual speech production is enacted by left ventro-lateral prefrontal cortex (vlPFC) supporting controlled access to conceptual representations (proactive control), and left mid-vlPFC supporting post-retrieval lexical selection (reactive control). In this functional magnetic resonance imaging (fMRI) study we demonstrate that multilingual word retrieval is supported by similar prefrontal mechanisms. However, differently from monolinguals, multilingual speakers retrieve words by applying proactive control on lexical representations to reduce cross-language interference. Alternatively to what it has been proposed by one of the most influential models, here we show that this proactive modulation is achieved by boosting the activation of lexical representations of the target language.

## 1. Introduction

Speech production is one of the most fundamental activities of humans. A core cognitive operation involved in this skill is word retrieval from the mental lexicon. Previous evidence has suggested that lexical-semantic knowledge is retrieved by left anterior vlPFC (proactive control), whereas instead task-relevant lexical representations are selected by left mid-vlPFC (reactive control) *via* suppression of lexical competitors (Badre et al., 2005, Crescentini et al., 2010; but see Snyder et al., 2011).

For multilingual individuals, comprising approximately half of the world’s population, this capacity includes management of two or more languages. Therefore, a critical question is whether multilingual lexical access is supported by the same prefrontal mechanisms as monolingual lexical access. One possibility is that multilingual word retrieval might involve not only reactive suppression of lexical competitors (Green, 1998; Green and Abutalebi, 2013), but also proactive down-regulation of non-target language. In fact, since preparation to speak in multilinguals involves getting prepared to use the target language, in some circumstances lexical representations of the non-target language might be inhibited even before knowing the specific words to be uttered.

This functional magnetic resonance imaging (fMRI) study investigated whether different vlPFC regions support proactive and reactive control processes in multilingual lexical access, and whether proactive control involves suppression of lexical representations. Hence, we tested multilingual speakers in a picture-naming task that required them to switch between their dominant and less-dominant language. As in previous studies (Czernochowski et al., 2015; see Ruge et al., 2013 for a review), we manipulated the cue-target interval (CTI) to either engage (long CTI) or prevent proactive control processes (short CTI). We also manipulated the cognate status of the pictures to-be-named (non-cognates *versus* cognates), to measure selective pre-activation of the target language (see below).

To test the two-process model (Badre and Wagner, 2007), we determined two regions of interest (ROIs), i.e., the left anterior and mid-vlPFCs, and we established *via* functional connectivity (FC) the networks related to these main ROIs during language switching. Firstly, we hypothesize that left anterior and mid-vlPFCs would support dissociable mechanisms. Hence, we expected them to be strongly co-activated with different brain regions. Particularly, we expected activation of left mid-vlPFC to show tighter coupling with activation in left inferior parietal lobe/supramarginal gyrus (IPL/SMG), reflecting attentional mechanisms for response conflict (Badre and Wagner, 2006). Instead, we expected activation of left anterior vlPFC to show stronger coupling with activation in left middle temporal gyrus (MTG), a region associated with lexical processing (Badre and Wagner, 2007; Strijkers et al., 2017).

To test our second hypothesis that left anterior vlPFC and mid-vlPFC would support proactive and reactive control processes respectively, we also examined neural responses in the two vlPFC ROIs and other ROIs determined from FC analysis. On one hand, we were expecting increased activation in anterior vlPFC for long *versus* short CTIs. On the other, we were expecting increased activation in mid-vlPFC for the opposite contrast, reflecting mostly reactive control (Badre and Wagner, 2007).

To test our third hypothesis that proactive control reduces co-activation of the two languages (preparation to speak in the target language), we examined the interaction between “cognate status” and CTI. The cognate status refers to two different classes of words that despite having the same meaning, they differ in the extent to which their phonological and orthographical representations are shared across languages. Hence, cognates are those translation words that have similar orthographic-phonological forms in the two languages of a bilingual (e.g., tomato—English, tomate—Spanish). Instead, non-cognates are those translation equivalents that share only their meaning in the two languages (e.g., apple—English, manzana— Spanish). Typically, cognate words as compared to non-cognate words are retrieved and processed faster (e.g., Christoffels et al., 2007). Importantly, behavioural and neural differences between non-cognate and cognate processing typically indicate that lexical representations of two languages are simultaneously active (cognate effect; Christoffels et al., 2007). Hence, in the present study we expected a reduced cognate effect during long *versus* short CTIs in the anterior vlPFC, but not in the mid-vlPFC (Badre and Wagner, 2007).

Finally, to test our fourth hypothesis that proactive control may involve language inhibition, we assessed the pattern of CTI and cognate status interaction. The inhibitory control model (ICM) (Green, 1998; Green and Abutalebi, 2013) proposes that lexical representations are controlled at multiple levels. One level of control is exerted locally by the “language task schemas” that regulate directly the outputs from the lexico-semantic system by selecting target lexical representations and by inhibiting non-target lexical representations. A second level of control is implemented by a supervisory attentional system (the SAS). The SAS proactively alters the activation level of a selected language task schema. This modulation might ultimately bias the activation of target language representations. Whilst SAS modulation on the selected language task schema might not swipe away completely the consequence of reactive inhibition, it may nevertheless reduce it. This may be achieved by down-regulating the activation level of the non-target language task schema. Hence, if proactive control involves inhibition of the non-target language (via SAS), this should may be observed for cognates when comparing long *versus* short CTIs. In fact, cognates (e.g., *tomato* in English) should benefit from co-activation of the translation equivalent in the non-target lexicon (e.g., *tomate* in Spanish) when both languages are co-activated (short CTI), and lose such benefit when the non-target lexicon is proactively inhibited (long CTI). In other words, neural activation in anterior vlPFC should vary between long and short CTI when naming cognates, but not non-cognates.

## 2. Materials and Methods

### 2.1. Participants

A total of 30 Spanish-Basque-English multilingual volunteers took part in the experiment. Four participants were excluded from further analyses due to excessive head motion during scanning (see *MRI data acquisition and analysis* section below). Moreover, a criterion for fMRI data inclusion in the analyses was adopted such as task blocks in which participants produced more than one erroneous response were modelled separately to exclude them from the main analyses. Importantly, given the present experiment was fMRI-blocked design, this criterion ensured to include only blocks containing at least 80% of correct responses. Thus, three additional participants were excluded because they had more than 23% of blocks with more than one error. The final study sample consisted of 23 participants (mean age = 24 years ± 4; 12 females).

For all the participants Spanish was the first and dominant language (L1), whereas English was a non-dominant language, acquired later in life (i.e., L3; mean age of L3 acquisition = 5 years ± 3). All participants were right-handed and had normal or corrected-to-normal vision. No participant had a history of major medical, neurological disorders, or treatment for a psychiatric disorder. The study protocol was approved by the Ethics Committee of the Basque Center for Cognition, Brain and Language (BCBL) and was carried out in accordance with the Code of Ethics of the World Medical Association (Declaration of Helsinki) for experiments involving humans. Prior to their inclusion in the study, all subjects provided informed written consent. Participants received monetary compensation for their participation.

### 2.2. Stimuli

Two-hundred and eight line drawings of common and concrete objects, belonging to a wide range of semantic categories (e.g., animals, body parts, buildings, furniture) were selected for the study (International Picture Naming Project, see Szekely et al., 2004). Fifty percent of the selected pictures (160 experimental and 48 filler pictures) were cognates and the remaining 50% were non-cognates. Experimental pictures were matched for visual complexity (reported in the IPNP database) [t (158) = 0.141, p = 0.888] and lexical frequency in Spanish and English [t(158) = -0.689, p = 0.492; and t(158) = -0.689, p = 0.73 respectively].

### 2.3. Experimental task and procedure

Participants were presented with a language-switching task divided in eight experimental runs. Our analyses focused on switching blocks that were intermixed with single-language naming blocks (L1 naming and L3 naming) during functional data collection. Within each switching block the two languages were continuously alternated (e.g., L1, L3, L1, L3, L1 or L3, L1, L3, L1, L3). We manipulated two variables: CTI (long, short) and the cognate status of the pictures (cognates, non-cognates). This resulted in a total of 16 switching blocks for each condition of interest (i.e., short cognates, long cognates, short non-cognates and long non-cognates). Each naming block included 5 experimental and two filler to-be-named pictures. Filler pictures had the same properties of the experimental pictures. Importantly, filler pictures as well as single-language naming blocks were added in order to reduce task predictability and they were modelled separately in the fMRI analyses. Therefore, a total of 16 block including 80 experimental pictures were included in each experimental condition of interest.

Before participants underwent MRI scanning, they received the instructions of the task, were familiarized with picture names in both languages and performed a practice session identical to the actual fMRI task. Instructions emphasized both speed and accuracy. During familiarization, the experimenter suggested the correct response when participants could not retrieve the name of the object depicted in the picture. This was done in order to reduce the likelihood of errors during the actual fMRI experiment. Participants were also instructed to minimize jaw–tongue movements while producing overt vocal responses to pictures and to say “skip” when they were not able to retrieve the name of the picture.

Once entered in the MRI scanner, participants were presented with written instruction again. Then the first trial started with a “language cue” (i.e., Spanish or English flag) presented during 100 ms and then followed by the target picture for 700 ms. During the time interval between the cue and the picture (i.e., CTI) a fixation cross was presented during either 50 ms or 900 ms. Hence, the total time between the cue and the target picture presentation was either 150 ms (i.e., short CTI) or 1000 ms (i.e., long CTI) respectively. Since every trial had a fixed duration (i.e., 3 s), the time between the presentation of the target picture and the beginning of the following trial was variable (either of 2850 ms or 2000 ms).

Four resting fixation baseline intervals were included within each functional run in which a fixation cross was displayed for 18 s at the center of the screen. Importantly, the task was presented with an fMRI blocked design, by means of Presentation software (Neurobehavioral systems: http://www.neurobs.com/). The choice of fMRI blocked design rather single trial event-related design is particularly related to maximise statistical power (Friston et al., 1999).

Vocal responses to each picture were classified as correct responses, incorrect responses and omissions (non-responses) for accuracy assessment. Finally, the background noise in the scanner did not allow obtaining accurate measures for naming latencies. Hence, behavioural analyses are reported only for accuracy (see “Behavioural data analyses”).

### 2.4. Behavioural data analyses

Behavioural analyses were performed on accuracy measures in order to explore the consequences of proactive (long *versus* short CTI) and reactive control (short *versus* long CTI) on multilingual lexical access (cognate effect). Therefore, we conducted a 2 (CTI: long, short) X 2 (cognate status: cognate, non-cognate) repeated measures ANOVA.

Importantly, we first excluded from this analysis those blocks that were not included in the fMRI analysis, i.e., all the blocks in which more than an erroneous response was found (10.1%, SD = 6.3 of the blocks in total). Then, productions of incorrect names and verbal disfluencies (stuttering, utterance repairs, and production of nonverbal sounds) were considered erroneous responses. Conversely, responses were considered correct whenever the expected name was given, but also when participants used consistently an appropriate category label for the item (e.g., synonyms) that did not affect its cognate status.

### 2.5. MRI data acquisition and analysis

Whole-brain MRI data acquisition was conducted on a 3-T Siemens TRIO whole-body MRI scanner (Siemens Medical Solutions) using a 32-channel whole-head coil. Snugly fitting headphones (MR Confon) were used to dampen background scanner noise and to enable communication with experimenters while in the scanner. Participants viewed stimuli back-projected onto a screen with a mirror mounted on the head coil. To limit head movement, the area between participants’ heads and the coil was padded with foam and participants were asked to remain as still as possible and to minimize jaw–tongue movements while producing vocal responses. Participants’ responses were recorded with a 40 dB noise-reducing microphone system (FOMRI-III, Optoacoustics Ltd). A dual adaptive filter system subtracted the reference input (MRI noise) from the source input (naming) and filtered the production instantly while recording the output. This optic fiber microphone was also mounted on the head coil and wired to the sound filter box, of which the output port was directly wired to the audio in-line plug of the computer sound card. The audio files were saved and analyzed to obtain participants’ in-scanner accuracy.

Functional images were acquired in 8 separate runs using a gradient-echo echo-planar pulse sequence with the following acquisition parameters: time repetition (TR)= 2500 ms, time echo (TE) = 25 ms, 43 contiguous 3-cubic mm axial slices, 0-mm inter-slice gap, flip angle = 90º, field of view (FoV) = 192 mm, 64 × 64 matrix, 235 volumes per run. Each functional run was preceded by 4 functional dummy scans to allow T1-equilibration effects that were discarded. High-resolution MPRAGE T_1_-weighted structural images were also collected for each participant with the following parameters: TR = 2300 ms, TE = 2.97 ms; flip angle = 9°, FoV = 256 mm, voxel size = 1-cubic mm, 150 slices.

#### Preprocessing

Standard SPM8 (Wellcome Department of Cognitive Neurology, London) preprocessing routines and analysis methods were employed. Images were corrected for differences in timing of slice acquisition and were realigned to the first volume by means of rigid-body motion transformation. Motion parameters extracted from the realignment were used, after a partial spatial smoothing of 4-mm full width at half-maximum (FWHM) isotropic Gaussian kernel, to inform additional motion correction algorithms implemented by the Artifact Repair toolbox (ArtRepair; Stanford Psychiatric Neuroimaging Laboratory) intended to repair outlier volumes with sudden scan-to-scan motion exceeding 0.5 mm and/or 1.3 % variation in global intensity, and that corrects these outlier volumes via linear interpolation between the nearest non-outliers time points (Mazaika et al., 2009). To further limit the influence of motion in our fMRI design, participants with more than 10% of to-be-corrected outlier volumes across functional runs were excluded. Before applying this additional motion correction procedure we also checked for participants who showed a drift over 3-mm/degrees in any of the translation (x, y, z) and rotation (yaw, pitch, roll) directions within each functional run. As a result of applying these motion correction criteria, we excluded a total of 3 participants from further data analyses.

After volume repair, structural and functional volumes were spatially normalized to T1 and echo-planar imaging templates, respectively. The normalization algorithm used a 12-parameter affine transformation together with a nonlinear transformation involving cosine basis functions. During normalization, the volumes were sampled to 3-mm cubic voxels. Templates were based on the MNI305 stereotaxic space (Cocosco et al., 1997), an approximation of Talairach space (Talairach and Tournoux, 1998). Functional volumes were then spatially smoothed with a 7-mm FWHM isotropic Gaussian kernel. Finally, time series were temporally filtered to eliminate contamination from slow drift of signals (high-pass filter: 128 s).

#### Whole-brain analysis

Statistical analyses were performed on individual participant data using the general linear model (GLM). The fMRI time series data were modelled by a series of impulses convolved with a canonical hemodynamic response function (HRF). The experimental conditions were modelled as 15 s epochs from the onset of the presentation of the first stimulus within each block until the end of the presentation of the last experimental stimulus within the block. The resulting functions were used as covariates in a GLM, along with the motion parameters for translation (i.e., x, y, z) and rotation (i.e., yaw, pitch, roll) as covariates of non-interest. The least-squares parameter estimates of the height of the best-fitting canonical HRF for each condition were used in pairwise contrasts. Contrast images, computed on a participant-by-participant basis, were submitted to group analyses. At the group level, whole-brain contrast for all switching conditions (Switch > Rest) was computed by performing a one sample t-test on these images, treating participants as a random effect. The standard statistical threshold for whole-brain maps was a voxel-wise corrected false discovery rate (FDR) set at q < 0.001. Brain coordinates throughout the manuscript are reported in MNI atlas space (Cocosco et al., 1997).

#### Seed-based whole-brain FC analysis

To identify the functional networks coupled with anterior vlPFC and mid-vlPFC activation during language switching, two separate seed-based whole-brain FC analyses were performed. The seeds used in these whole-brain FC analyses were identified from the Switch > Rest whole-brain functional t-contrast across all participants (thresholded at q < 0.001 FDR voxel-wised corrected for multiple comparisons). FC analysis was conducted *via* the beta-series correlation method (Rissman et al., 2004), implemented in SPM with custom Matlab scripts. The beta series correlation is an established FC method which presents a series of advantages that are relevant for our study, compared to other voxel-wise analyses of connectivity. It is also a highly appropriate FC method for slow event-related designs, which is akin to the present design wherein the different conditions were presented in epochs lasting 15 seconds.

The canonical HRF in SPM was fit to each trial from of each experimental condition and the resulting parameter estimates (i.e., beta values) were sorted according to the study conditions to produce a condition-specific beta series for each voxel. The beta series associated with these seeds were correlated with voxels across the entire brain to produce beta correlation images for each subject for the contrast Switch > Rest. These contrasts were subjected to an arc-hyperbolic tangent transform (Fisher, 1921) to allow for statistical inference based on the correlation magnitudes. Group-level one sample t-tests FC maps were performed on the resulting subject Switch > Rest contrast images for each of the selected seeds (i.e., left anterior vlPFC and left mid-vlPFC) using a voxel-wise corrected Family Wise Error (FWE) threshold of p < 0.05. Given our hypothesis in regard to the involvement of the anterior vlPFC in proactive control and mid-vlPFC in reactive control during language switching, we expected to observe two distinct functional networks associated with each of these seed-based whole-brain FC analyses. Hence, to determine differential coupling strength between the anterior vlPFC and mid-vlPFC networks, these maps were submitted to a paired t-test, using a voxel-wise FDR correction set at q < 0.05.

#### ROI analysis

ROI analyses were conducted on the regions reported in **Table 1** to examine interactions between CTI and cognate status for regions of interest (e.g., Badre and Wagner, 2006; Badre and Wagner, 2007) within the seed-based FC networks associated with proactive (i.e., anterior vlPFC) and reactive control (i.e., mid-vlPFC). Note that identifying the ROIs from the networks derived from seed-based whole-brain FC during language switching (i.e., Switch > Rest) avoid biases associated with the CTI and cognate effects, only constraining the ROIs to voxels that were coupled with left anterior and mid-vlPFC across our experimental fMRI design conditions. Importantly, in this analysis, as well as in pairwise FC analysis (see below), we employed 5-mm radius ROIs spheres centered at the highest local maximas within each ROI, also for left anterior vlPFC and left mid-vlPFC regions. This was done to ensure that differences in coupling strength in pairwise FC analysis (see below) were not determined by differences in the sizes of the functionally defined ROIs.

**Table 1.**
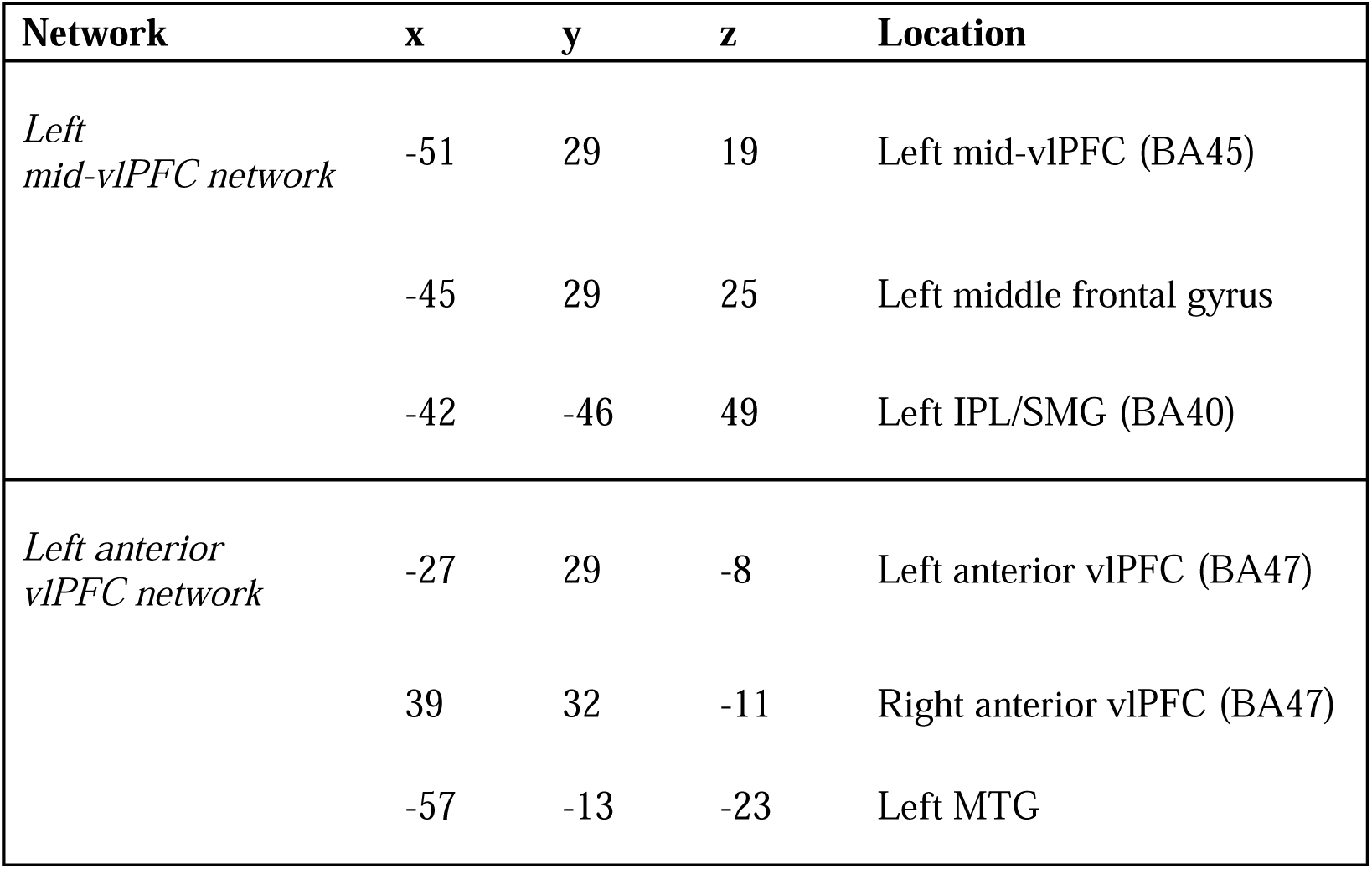
Coordinates of ROIs (spheres) for each functional connectivity network.

Parameter estimates (i.e., beta values) for each ROI were extracted with the MARSBAR toolbox (Brett et al., 2002). Then, to specifically examine to what extent multilingual lexical access was affected by preparatory processes, we submitted percent signal change values from each ROI to a 2 (CTI: long, short) X 2 (Cognate status: cognate, non-cognate) repeated measures ANOVAs. Bonferroni corrections for multiple comparisons were applied to the post-hoc analyses.

#### Pairwise FC analysis

Finally, to examine whether coupling strength between pairs of ROIs within these two networks was modulated by CTI variables and/or the cognate status, pairwise FC analysis were conducted using the beta-series correlation method (Rissman et al., 2004). The canonical HRF in SPM was fitted to each occurrence of each condition and the resulting parameter estimates (i.e., beta values) were sorted according to the study conditions to produce a condition-specific beta series for each voxel. To examine pairwise functional connectivity between the ROIs, beta correlation values for each pair of ROIs per subject and condition were then calculated. Then, an arc-hyperbolic tangent transform (Fisher, 1921) was applied at the subject level to the beta-series correlation values (*r* values) of each pair of ROIs and each study condition. Since the correlation coefficient is inherently restricted to range from -1 to 1, this transformation served to make its null hypothesis sampling distribution approach that of the normal distribution. Then, in order to test for significant differences in coupling strength between conditions of interest, we submitted these fisher’s z normally distributed values for each pair of ROIs, participant and condition, to paired t-tests using a FDR correction for multiple comparisons set at q < 0.05.

## 3. Results

### 3.1. Behavioural results

We performed behavioural analyses on accuracy measures to explore the behavioural consequences of proactive and reactive control during multilingual lexical access. Results revealed a significant main effect of CTI [F (1, 22) =11.438, p = 0.003, *η*p^2^ = 0.342], showing that responses for short CTI (93.9%, SD = 1) were more accurate than those for long CTI (92.2%, SD = 3) (see **Figure 1**). Results also revealed more accurate responses for cognates (94.5%, SD = 2) as compared to non-cognates (91.6%, SD = 3) [main effect of cognate status: F (1, 22) = 33.152, p < 0.001, *η*p^2^ = 0.601].

**Figure 1.**
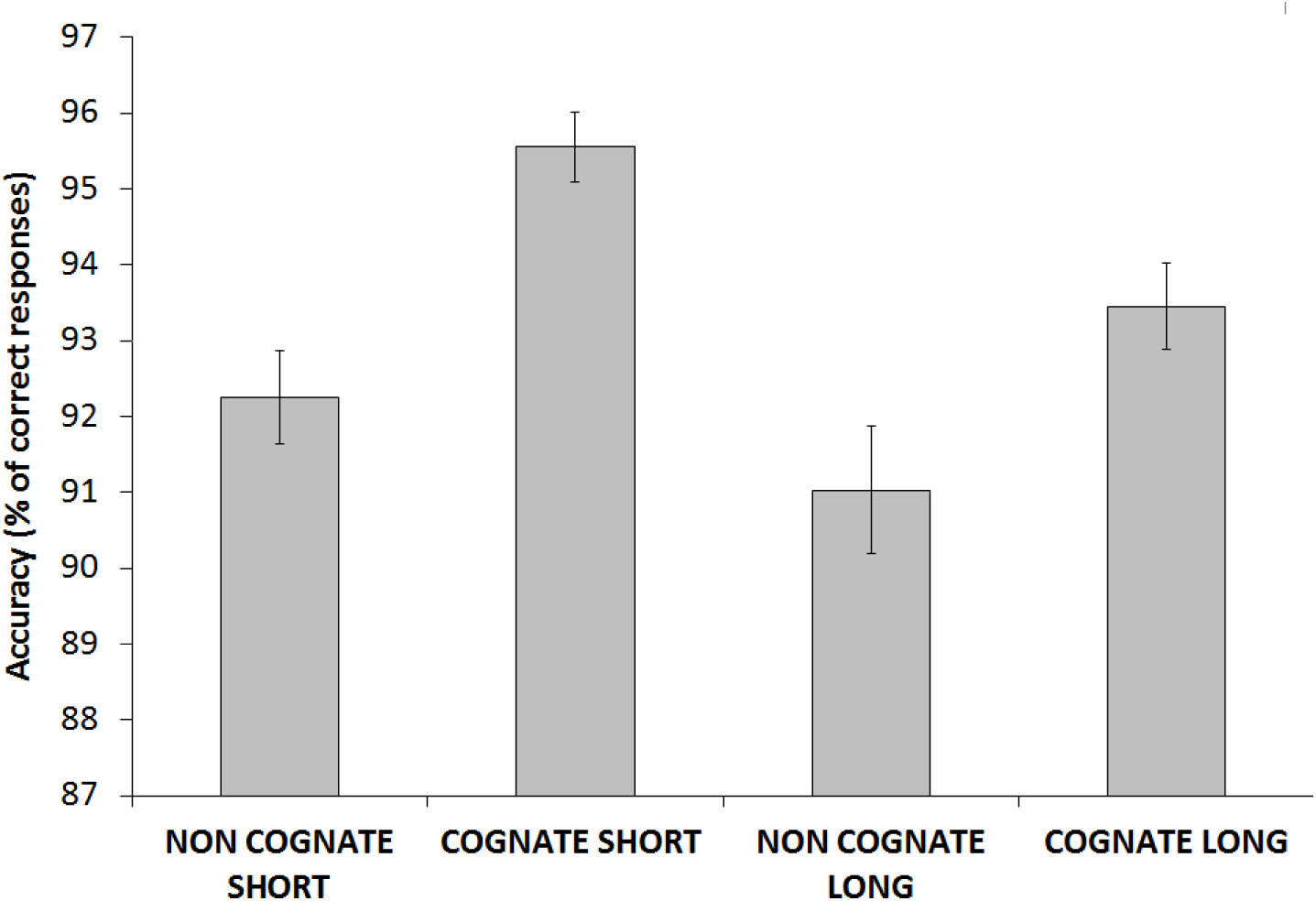
Behavioural results for accuracy. Error bars denote standard errors (SEs).

The interaction between cognate status and CTI was not significant [F (1, 22) = 1.035, p = 0.32, *η*p^2^ = 0.045], suggesting that the cognates and non-cognates were similarly modulated by long and short CTIs.

### 3.2. fMRI results

#### Whole-brain contrast

To identify brain regions associated with language switching across all participants, we computed a whole-brain T-contrast for Switch > Rest. The contrast revealed the involvement of a bilateral network of regions, including both language control and representational areas. Importantly, both left anterior and mid-vlPFCs were significantly activated by this contrast (see **Figure 2**).

**Figure 2.**
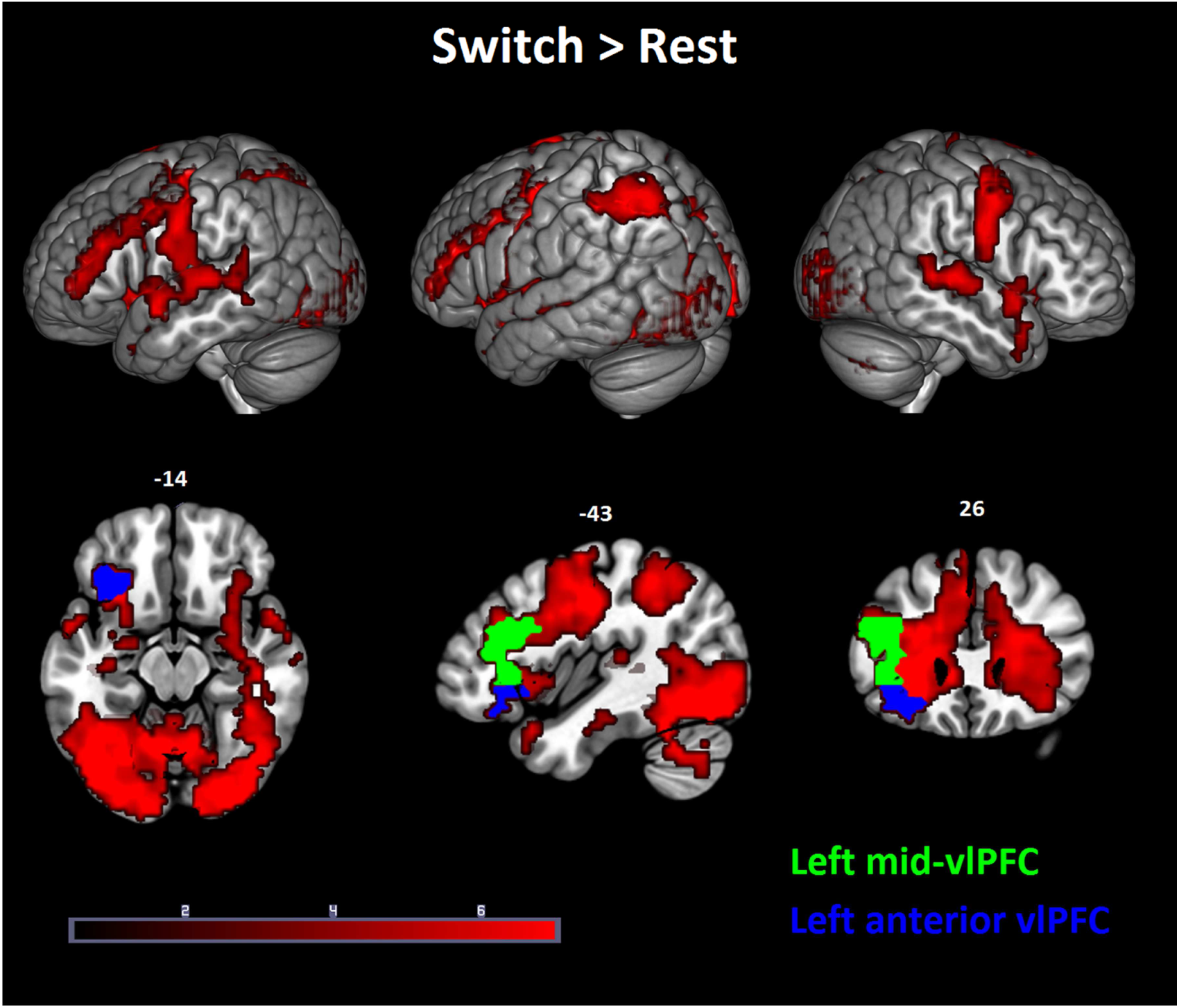
Whole-brain Switch > Rest contrast. Seed-regions used in the subsequent whole-brain FC analyses are highlighted in green and blue colours respectively for left mid-vlPFC and left anterior vlPFC.

#### Seed-based whole-brain FC results

This analysis was aimed at identifying which areas were strongly coupled with left mid-vlPFC and left anterior vlPFC during multilingual word retrieval. Whole-brain FC from left mid-vlPFC (left: -43, 28, 15; 10840 mm^3^) and left anterior vlPFC (left: -35, 28, -10; 4392 mm^3^) (see green and blue regions in **Figure 2**) revealed partially overlapping brain networks, including both cortical and subcortical cognitive control regions, as well as temporal brain areas (see **Figure 3a**).

**Figure 3.**
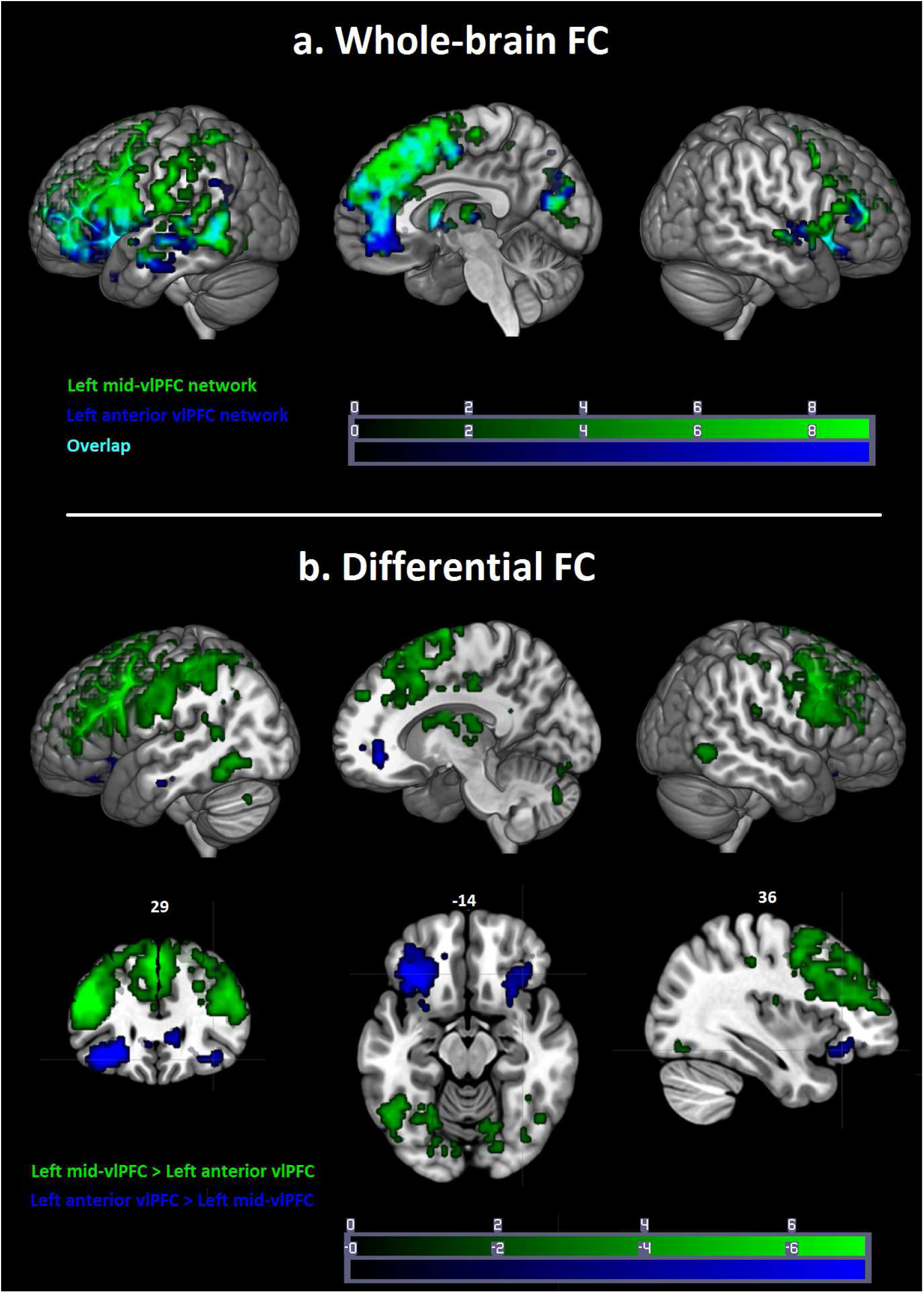
a) Whole-brain FC for left mid-vlPFC (green) and left anterior vlPFC (blue) networks; b) Differential FC for left mid-vlPFC network > left anterior vlPFC network (green) and for left anterior vlPFC network > left mid-vlPFC network (blue).

Paired t-test results indicated differential coupling strength between whole-brain FC originating from these two seeds. On one hand, whole-brain FC from left mid-vlPFC *versus* left anterior vlPFC was significantly tighter in lateral dorsal PFC regions, left IPL/SMG, and posterior temporal regions. On the other hand, whole-brain FC from left anterior vlPFC *versus* left mid-vlPFC was significantly stronger in left MTG and right anterior vlPFC (see **Figure 3b**).

#### ROI results

We conducted ROI analyses to examine interactions between CTI and cognate status variables. The results (see **Table 2 and Figure 4**) can be summarised as follows: The left IPL/SMG and the two ROIs within the mid-vlPFC network were not sensitive to the CTI manipulation, suggesting that they were similarly recruited for reactive and proactive control. Moreover, these regions were sensitive to the cognate manipulation and showed increased neural responses for cognates as compared to non-cognates (i.e., cognate status main effect). This effect was also qualified by the significant interaction between cognate status and CTI. Follow-up t-tests revealed that the significant interaction was determined by larger cognate effect (cognate *versus* non-cognate difference) during short as compared to long CTIs [left mid-vlPFC (BA45): t (22) = 2.133, p=0.044; left Middle Frontal Gyrus: t (22) = 3.056, p=0.006; left IPL/SMG: t (22) = 4.493, p<0.001]. The left MTG showed sensitivity to the cognate status manipulation with increased activation for cognates and reduced activation for non-cognates.

**Table 2.** Summary of results of ROI analyses.

| Region of interest | CTI (Long > Short) | Cognate status (Cognate > Non cognate) | Interaction | Post hoc test (Bonferroni corrected) | Follow up t-tests |
| --- | --- | --- | --- | --- | --- |
| 1. Left mid-vIPFC (BA45) | F (1, 22) = 1.46<br>$p = 0.706$<br>$\eta^2 = 0.007$ | F (1, 22) = 16.584<br>$p = 0.001$<br>$\eta^2 = 0.430$ | F (1, 22) = 4.546<br>$p = 0.044$<br>$\eta^2 = 0.171$ | a) Short CTI: Cognate > Non cognate ($p < 0.001$)<br>b) Long CTI: Cognate > Non cognate ($p = 0.009$) | $a > b$ [t (22) = 2.133, $p = 0.044$] |
| 2. Left Middle Frontal Gyrus | F (1, 22) = 0.202<br>$p = 0.658$<br>$\eta^2 = 0.009$ | F (1, 22) = 16.46<br>$p = 0.001$<br>$\eta^2 = 0.428$ | F (1, 22) = 11.6<br>$p = 0.003$<br>$\eta^2 = 0.345$ | a) Short CTI: Cognate > Non cognate ($p < 0.001$)<br>b) Long CTI: Cognate > Non cognate ($p = 0.028$) | $a > b$ [t (22) = 3.056, $p = 0.006$] |
| 3. Left IPL/SMG (BA40) | F (1, 22) = 0.014<br>$p = 0.908$<br>$\eta^2 = 0.001$ | F (1, 22) = 24.899<br>$p < 0.001$<br>$\eta^2 = 0.531$ | F (1, 22) = 23.214<br>$p < 0.001$<br>$\eta^2 = 0.513$ | a) Short CTI: Cognate > Non cognate ($p < 0.001$)<br>b) Long CTI: Cognate > Non cognate ($p = 0.002$) | $a > b$ [t (22) = 4.493, $p < 0.001$] |
| 4. Left anterior vIPFC (BA47)* | F (1, 21) = 5.922<br>$p = 0.024$<br>$\eta^2 = 0.220$ | F (1, 21) = 46.459<br>$p < 0.001$<br>$\eta^2 = 0.689$ | F (1, 21) = 6.003<br>$p = 0.023$<br>$\eta^2 = 0.222$ | Non cognate : Long CTI > Short CTI ($p = 0.008$)<br>a) Short CTI: Cognate > Non cognate ($p < 0.001$)<br>b) Long CTI: Cognate > Non cognate ($p < 0.001$) | $a > b$ [t (21) = 4.493, $p < 0.001$] |
| 5. Right anterior vIPFC (BA47) | F (1, 22) = 0.775<br>$p = 0.388$<br>$\eta^2 = 0.034$ | F (1, 22) = 9.109<br>$p = 0.006$<br>$\eta^2 = 0.293$ | F (1, 22) = 16.883<br>$p < 0.001$<br>$\eta^2 = 0.434$ | Non cognate : Long CTI > Short CTI ($p = 0.029$)<br>Short CTI: Cognate > Non cognate ($p = 0.001$) | |
| 6. Left MTG | F (1, 22) = 0.008<br>$p = 0.927$<br>$\eta^2 < 0.001$ | F (1, 22) = 9.875<br>$p = 0.005$<br>$\eta^2 = 0.31$ | F (1, 22) = 0.257<br>$p = 0.617$<br>$\eta^2 = 0.012$ | | |

\* = Beta values above the threshold was found only for 21 participants.

**Figure 4.**
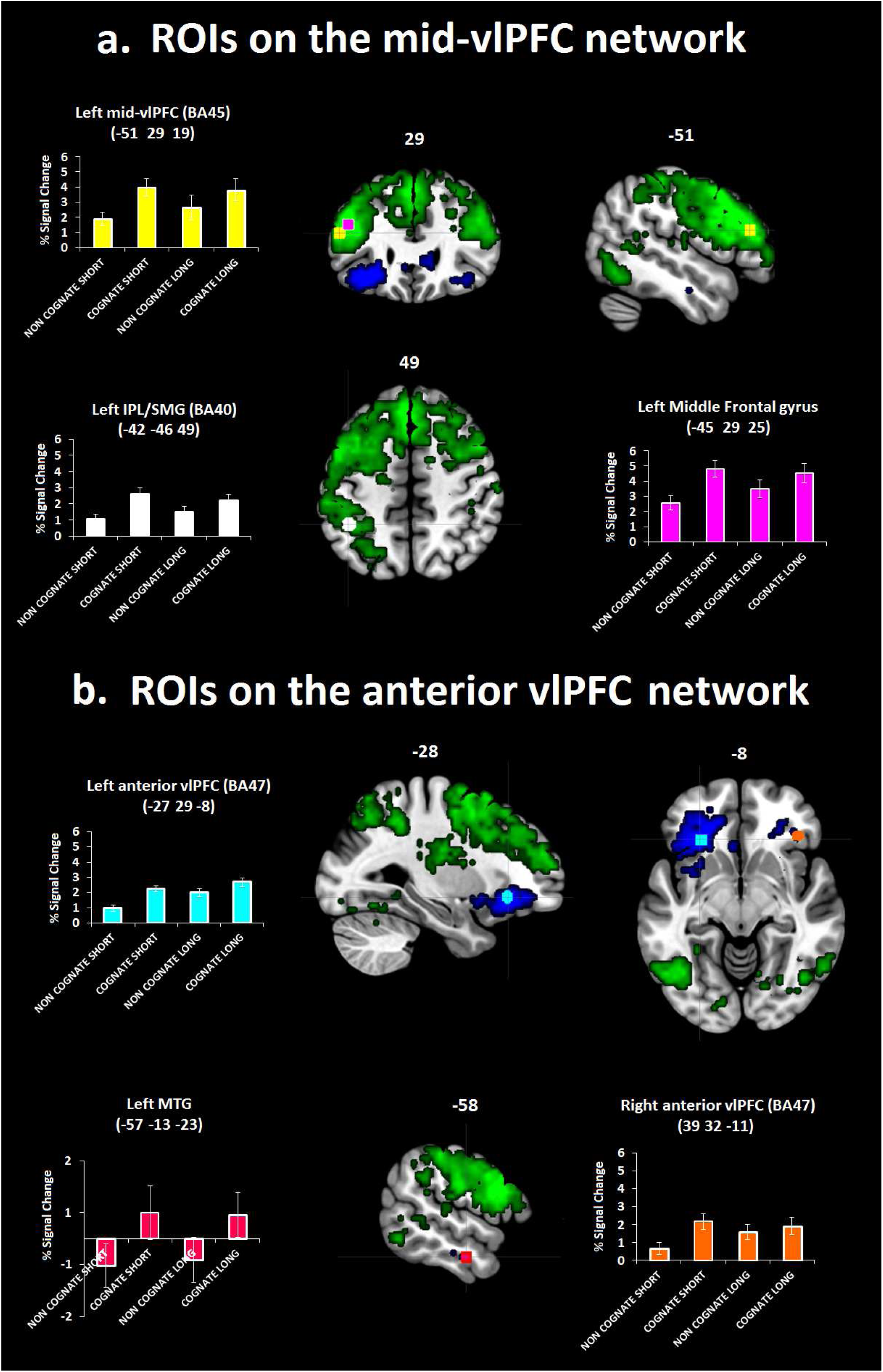
ROI analyses for regions (a) within left mid-vlPFC network, including left mid-vlPFC, left MFG and left IPL/SMG; and (b) within left anterior vlPFC network, including left and right anterior vlPFC and left MTG. Brain coordinates correspond to the MNI coordinates for the center of mass of each ROI.

The left and right anterior vlPFCs were both sensitive to cognate status manipulation. Moreover, in these areas, a significant interaction between CTI and cognate status revealed that activation for non-cognates was increased during long as compared to short CTIs (proactive modulation). Consequently, the difference between cognates and non-cognates was reduced in left anterior vlPFC [t (21) = 2.45, p=0.023] during long *versus* short CTIs and eliminated in right anterior vlPFC.

#### Pairwise FC results

We further investigated whether coupling strength between our *a priori* regions of interest could be modulated by CTI and cognate status conditions. Hence, we conducted pairwise FC analysis between the selected ROIs.

FC between pairs of ROIs was modulated by proactive control (long *versus* short CTIs) only for non-cognates. Precisely, increased FC was observed for long *versus* short CTIs between left anterior vlPFC (BA47) and left mid-vlPFC (BA45) (t = 2.34, q < 0.05), and between right anterior vlPFC (BA47) and left mid-vlPFC (BA45) (t = 2.58, q < 0.05) (see **Figure 5a**). Accordingly, when preparatory processes were involved (long CTIs), stronger coupling for non-cognates *versus* cognates was observed between left anterior vlPFC (BA47) and left mid-vlPFC (BA45) (t = 2.47, q < 0.05), and between right anterior vlPFC (BA47) and left IPL/SMG (t = 2.21, q < 0.05) (see **Figure 5b)**.

**Figure 5.**
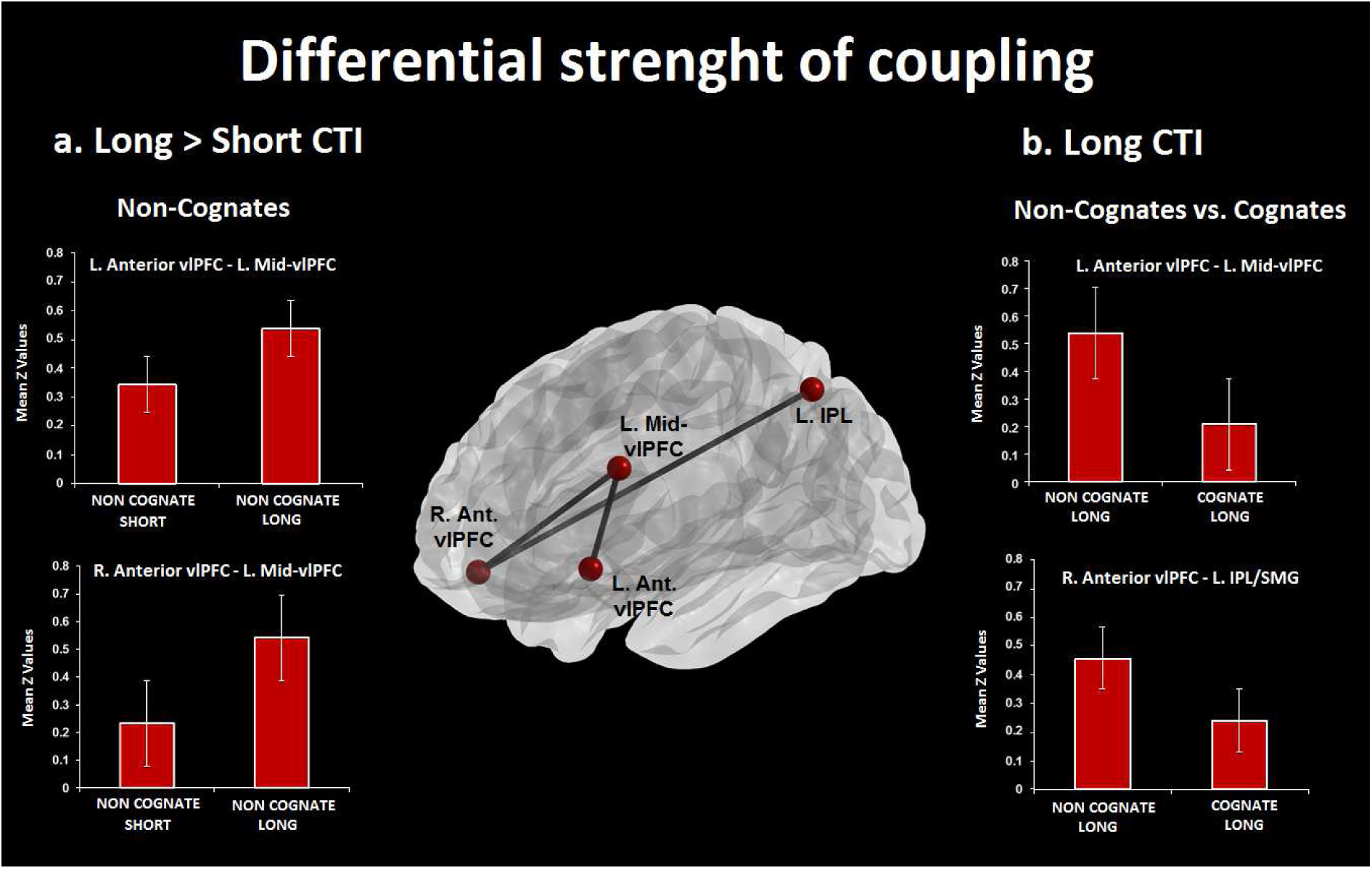
Pairwise FC results among ROIs showing differential strength of coupling for a) long > short CTI and b) long CTI. Error bars refer to SE.

In summary, we observed that the strength of coupling between different areas was particularly modulated by proactive control (long > short CTI). Importantly, in accord with ROI results (see above) the observed proactive modulation seemed to affect particularly non-cognates (long: non-cognates > cognates) that showed increased coupling between right vlPFC and left IPL/SMG.

## 4. Discussion

This fMRI study addressed whether word retrieval in multilingual speakers is supported by dissociable vlPFC mechanisms reflecting proactive and reactive control processes, and whether multilinguals use proactive control to suppress lexical representations of the non-target language. Our findings support the two-process model of lexical-semantic control (Badre and Wagner, 2007) and extend it for the first time to multilingual lexical access. By employing FC we were able to reveal a clear functional segregation between functional networks related to mid-vlPFC and anterior vlPFC, supporting our first hypothesis that these two regions enact dissociable mechanisms for reactive and proactive control, respectively (Badre and Wagner, 2007). Activation in left mid-vlPFC was coupled with activation in IPL/SMG. Instead, activation in left anterior vlPFC was coupled with activation in left MTG. Increased FC between left mid-vlPFC and left IPL/SMG is consistent with evidence showing that these areas are both engaged during reactive control in switching tasks (Badre and Wagner, 2006; Green and Abutalebi, 2013). The left IPL/SMG, a key region for phonological control (Hartwigsen et al., 2010), may support language selection enacted by left mid-vlPFC by biasing selection away from non-target phonological representations (Branzi et al., 2016; Abutalebi and Green 2016). Instead, increased FC between left anterior vlPFC and left MTG might reflect controlled retrieval of target lexical representations in the temporal lobe (Badre and Wagner, 2007).

In line with our second hypothesis that left anterior vlPFC would be recruited for proactive control specifically, ROI analyses revealed increased neural activation in this area for long *versus* short CTIs. Contrary to our predictions, however, left mid-vlPFC did not show the expected increased neural responses for short *versus* long CTIs. Indeed, this region was not sensitive to the CTI manipulation, suggesting a similar involvement for reactive and proactive control. It is possible that, when conditions allow doing so, language control relies particularly on proactive processes (Martin et al., 2016; Molnar et al., 2015). In language switching this strategy may be crucial to adjust performance according to continuously changing goals. Hence, this might result in a more extensive use of control areas during proactive control in general, that may rule out “specific” effects for reactive control (i.e., differential neural activation for short *versus* long CTI conditions). These results suggest that word retrieval in multilingual speakers is enacted by two distinct vlPFC areas, showing a different profile of regional engagement and network coactivation. The left mid-vlPFC supports language selection during proactive and reactive control and its coupling with left IPL/SMG might reflect phonological control. The left anterior vlPFC supports proactive control specifically and its coactivation with left MTG might reflect controlled access to lexical representations. Importantly, the ROIs within each network showed the same profile of engagement, with the only exception of MTG, an area whose profile of activation is consistent with a representational rather than a control role (see Badre and Wagner, 2007).

In line with our third hypothesis that proactive modulation in left anterior vlPFC, but not in mid-vlPFC, would affect language co-activation (i.e., cognate effect), during long *versus* short CTIs we observed both a reduction and elimination of the cognate effect in left and right anterior vlPFCs respectively. Nevertheless, and contrary to our prediction, follow-up t-tests revealed a reduced cognate effect also in left mid-vlPFC for long *versus* short CTIs. Despite this result may suggest that some proactive modulation is involved, it is not clear where this modulation may be exerted, since neural responses in mid-vlPFC for both cognates and non-cognates were not statistically different between long and short CTIs. Furtheremore, the fact that activation in left mid-vlPFC was not sensitive to the CTI manipulation further suggests that this area may not play a key role in proactive control. Taken together these results provide evidence that proactive control is enacted by the recruitment of bilateral anterior vlPFC for selection of target lexical representations. Accordingly, Wu and Thierry (2017) have recently shown that preparatory processes affect bilingual language selection. However, given that variables related to lexicalisation processes were not manipulated in that study, it is unclear whether preparatory processes were affecting lexical representations. Indeed, since processing a language-cue is preverbal (speakers do not know yet what they will say, but only which language to use), preparation might involve only a general task-schema (“to name in a given language”), without necessarily inducing any modulation on language-specific representations. Our study allows making this inference. In fact, by manipulating the cognate status of the to-be-named pictures we were able to assess how proactive control modulated neural responses for cognates and non-cognates, and therefore to elucidate the mechanisms underlying the reduction of language coactivation.

Our fourth main goal was to test whether proactive control may consist in inhibitory modulation of the non-target language. If so, we were expecting this effect to be observed for cognates when comparing long *versus* short CTIs. Precisely, we expected to observe neural activation in anterior vlPFC to vary between long and short CTIs when naming cognates. Contrary to our prediction, ROI results revealed that the proactive modulation in the anterior vlPFC was exerted only on non-cognates. This modulation consisted of increased neural responses for long *versus* short CTI conditions. Instead, proactive control did not modulate neural activation for cognates. In line with ROI results, our data also revealed that proactive control modulated the coupling strength between different areas, particularly for non-cognate representations. The stronger coupling for non-cognates as compared to cognates was observed between regions of the two networks, such as between left anterior vlPFC and left mid-vlPFC, and between right vlPFC and left IPL/SMG. The observed tighter coupling between areas from the two networks during multilingual language production is an interesting and not previously reported finding. One possibility is that during long CTIs proactive control enacted by the anterior vlPFC is applied on lexical representations that are less activated (i.e., non-cognates) via interactions with brain areas involved in phonological control (IPL/SMG) and response selection (left mid-vlPFC).

The present findings provide important insights regarding multilingual language control mechanisms. First, the dissociation between cognates and non-cognates suggests that proactive control may be differently applied on the lexical representations with different phonological overlap. This observation is not in accord with the models that propose that language control is applied globally on the non-target language (Green and Abutalebi, 2013; Green, 1998). According to these proposals we should have observed cognates to be modulated by proactive processes to some extent. However, we did not observe such result across the different functional neuroimaging analyses here performed.

Second, contrary to what we hypothesised, our findings suggest that cognate representations are maintained active during the entire task, and those of non-cognates are selectively activated, rather than inhibited, *via* proactive control. Note that according to what we observed in behavioural results, it is hard to explain the increase of neural activity for non-cognates in long *versus* short intervals as reflecting increased retrieval demands for non-cognates.

The fact that proactive control reflects target language activation, would predict that the non-target language would still be activated despite proactive control. Ultimately, this activation would remain still ‘‘traceable” in the behavioral cognate effect in the long CTI condition. This interpretation and the results that we provide here are in accord with other non-inhibitory models of multilingual language control (Costa and Caramazza, 1999; Costa et al., 1999; Runnqvist et al., 2012). Finally, despite these results are not necessarily inconsistent with the ICM (Green and Abutalebi, 2013), since reactive inhibition may still occur (language task schema level), the fact we have not revealed any neural difference between short *versus* long intervals suggests that whereas proactive control can be applied, this may reduce the deployment of reactive (inhibitory) control processes.

## Conclusion

This study demonstrates for the first time that word retrieval in multilinguals is enacted by different portions of the vlPFC associated with proactive and reactive control processes and that multilinguals engage proactive control to reduce language co-activation. We report for the first time that this proactive modulation is achieved *via* activation of target lexical representations. These findings may also have an impact on the research on multilingual advantage in domain-general executive control (Antón et al., 2014; Branzi et al., 2018), as they suggest that the extent to which the advantage in proactive control could be observed might depend on the ratio between cognates and non-cognates in the languages of a multilingual.

## Acknowledgements

This work was supported by one grant from the European Research Council under the European Community’s Seventh Framework (FP7/2007–2013 Cooperation grant agreement 613465-AThEME). Francesca M. Branzi was partially supported by a postdoctoral fellowship from the European Union’s Horizon 2020 research and innovation programme, under the Marie Sklodowska-Curie grant agreement No 658341. Pedro M. Paz-Alonso was supported by grants (RYC-2014–15440, PSI2015-65696) from the Spanish Ministry of Economy and Competitiveness (MINECO), and a grant (PI2016-12) from the Basque Government. Manuel Carreiras was partially supported by grant ERC-2011-ADG-295362 from the European Research Council, and grant PSI2015-67353-R from the MINECO. BCBL acknowledges funding from Ayuda Centro de Excelencia Severo Ochoa SEV-2015–0490 from the MINECO

## References

Abutalebi J, Green DW. 2016. Neuroimaging of language control in bilinguals: neural adaptation and reserve. Biling Lang Cogn. 19:689–698.

Antón E, Duñabeitia JA, Estévez A, Hernández JA, Castillo A, Fuentes LJ, Davidson D, Carreiras M. 2014. Is there a bilingual advantage in the ANT task? Evidence from children. Front Psychol 5.

Badre D, Poldrack RA, Pare-Blagoev EJ, Insler RZ, Wagner AD. 2005. Dissociable controlled retrieval and generalized selection mechanisms in ventrolateral prefrontal cortex. Neuron. 47:907–918.

Badre D, Wagner AD. 2006. Computational and neurobiological mechanisms underlying cognitive flexibility. Proc Natl Acad Sci USA. 103:7186–7191.

Badre D, Wagner AD. 2007. Left ventrolateral prefrontal cortex and the cognitive control of memory. Neuropsychologia. 45:2883–901.

Branzi FM, Della Rosa PA, Canini M, Costa A, Abutalebi J. 2016. Language control in bilinguals: monitoring and response selection. Cereb Cortex. 26:2367–2380.

Branzi FM, Calabria M, Gade M, Fuentes LJ, Costa A. 2018. On the bilingualism effect in task switching. Biling Lang Cogn. 21:195–208.

Brett M, Anton JL, Valabregue R, Poline JB. 2002. Region of interest analysis using the MarsBar toolbox for SPM 9. NeuroImage 16:S497.

Christoffels IK, Firk C, Schiller NO. 2007. Bilingual language control: An event-related brain potential study. Brain Res. 1147:192–208.

Cocosco CA, Kollokian V, Kwan RKS, Pike GB, Evans AC. 1997. Brainweb: Online interface to a 3D MRI simulated brain database. NeuroImage. 5:S425.

Costa A, Caramazza A. 1999. Is lexical selection in bilingual speech production language-specific? Further evidence from Spanish–English and English–Spanish bilinguals. Biling Lang Cogn. 2:231–244.

Costa A, Miozzo M, Caramazza A. 1999. Lexical selection in bilinguals: Do words in the bilingual’s two lexicons compete for selection? J Mem Lang. 41:365–397.

Crescentini C, Shallice T, Macaluso E. 2010. Item retrieval and competition in noun and verb generation: an FMRI study. J Cogn Neurosci. 22:1140–1157.

Czernochowski D. 2015. ERPs dissociate proactive and reactive control: Evidence from a task-switching paradigm with informative and uninformative cues. Cogn Aff Behav Neurosci. 5:117–31.

Fisher RA. 1921. On the probable error of a coefficient of correlation deduced from a small sample. Metron. 1:3–32.

Friston KJ, Zarahn E, Josephs O, Henson RN, Dale AM. 1999. Stochastic designs in event-related fMRI. NeuroImage. 10:607–619.

Green D, Abutalebi J. 2013. Language control in bilinguals: The adaptive control hypothesis. J Cog Psych. 25:515–530.

Hartwigsen G, Baumgaertner A, Price CJ, Koehnke M, Ulmer S, Siebner HR. 2010. Phonological decisions require both the left and right supramarginal gyri. Proc Natl Acad Sci USA. 107:16494–16499.

Martin CD, Molnar M, Carreiras M. 2016. The proactive bilingual brain: using interlocutor identity to generate predictions for language processing. Sci Rep. 6:26171.

Mazaika PK, Hoeft F, Glover GH, Reiss AL. 2009. Methods and software for fMRI analysis of clinical subjects. NeuroImage. 47:S58.

Molnar M, Ibáñez-Molina A, Carreiras M. 2015. Interlocutor identity affects language activation in bilinguals. J Mem Lang. 81:91–104.

Rissman J, Gazzaley A, D’esposito M. 2004. Measuring functional connectivity during distinct stages of a cognitive task. NeuroImage. 23:752–763.

Ruge H, Brass M, Koch I, Rubin O, Meiran N, Von Cramon DY. 2005. Advance preparation and stimulus-induced interference in cued task switching: further insights from BOLD fMRI. Neuropsychologia. 43:340–355.

Runnqvist E, Strijkers K, Alario FX, Costa A. 2012. Cumulative semantic interference is blind to language: Implications for models of bilingual speech production. J Mem Lang. 66:850–869.

Snyder HR, Banich MT, Munakata Y. 2011. Choosing our words: retrieval and selection processes recruit shared neural substrates in left ventrolateral prefrontal cortex. J Cogn Neurosci. 23:3470–3482.

Strijkers K, Costa A, Pulvermüller F. 2017. The cortical dynamics of speaking: Lexical and phonological knowledge simultaneously recruit the frontal and temporal cortex within 200 ms. NeuroImage. 163:206–219.

Szekely A, Jacobsen T, D’Amico S, Devescovi A, Andonova E, Herron D, … Federmeier K. 2004. A new on-line resource for psycholinguistic studies. J Mem Lang. 51:247–250.

Talairach J, Tournoux P. 1988. Co-planar stereotaxic atlas of the human brain. New York: Thieme.

Wu YJ, Thierry G. 2017. Brain potentials predict language selection before speech onset in bilinguals. Brain Lang. 171:23–30.

